# Evaluation of Genetic Diversity in Cultivated and Exotic Germplasm Sources of Faba Bean Using Important Morphological Traits

**DOI:** 10.1101/2020.01.24.918284

**Authors:** Praveen Kumar, Prashant Kaushik

## Abstract

**Background and Objective:** Faba bean is an important crop for achieving nutritional food security, but there is very limited diversity in the cultivated varieties of faba bean. Moreover, genetic diversity is vital for its use in faba bean genetic imporvement.

**Material and Methods:** Here we determined the diversity in the sixty-four genotypes of faba bean of different agro-ecological origins. Plants were grown in randomized block design in three replications. Further, the genotypes were characterized based on the ten morphological traits.

**Results:** Highly significant differences were determined for all of the studied traits. Whereas, the number of cluster per plant was positively correlated with the pods per plants. Moreover, the trait number of cluster per plant determined the most substantial positive effect on seed yield.

**Conclusions:** Overall, our results indicate a wide range of variability for further selection and improvement of faba bean ideotype.

## Introduction

Pulses are well-known for their ability in maintaining soil fertility via nitrogen fixation, and therefore, pulses contribute significantly to the sustainability of the farming systems^1,2^. Although the pulses production and productivity are not much satisfactory to meet the current requirement of pulses. Moreover, the increase in pulses production could not keep pace with the demand of the increasing population^3,4^. Therefore, the development of high yielding varieties of nutritionally rich pulses can achieve higher production and sustainability^5^. In this direction, faba bean has excellent potential for yield and can bridge up the gap between production demand of pulses. Faba bean (*Vicia faba* L.) is also known as the broad bean, horse bean, pigeon bean, field bean, and bell bean is a major grain legume crop with a diploid chromosome number 2n=12^6^.

It is partially allogamous crop, grown for high seed protein content and superior biomass, ranking as the fourth most internationally important temperate crop after peas, chickpeas and lentils^7^. Faba bean is a multipurpose legume and an excellent resource of proteins, starch, cellulose and minerals^8,9^. Although, among the most commonly cultivated crops across the Indian subcontinent, faba bean has not only the highest crude protein content but also has the highest yield of protein per hectare^10^.

The diversity for the important traits is vital for the breeding program to keep on with changing the market demands^11,12^. The genetic resources in the form of landraces and crop wild relatives have played a significant role in providing useful traits to the modern cultivated varieties of a faba bean^7^. Moreover, the availability of adequate genetic variability, knowledge of criteria for screening and selection of desirable genotype are the prerequisites for the planning of a breeding programme for the development of ideal varieties. For active and purposeful exploitation of the genetic variability, it is necessary to recognize and measure such variability to utilize it in the breeding programme^13^. Thus, the following objectives were undertaken:

- To estimate the components of variability in different faba bean genotypes.
- To determine the extent of correlation between yield and yield component traits.
- To study the genetic divergence in different faba bean genotypes.

## Material and Methods

### Experimental Design and Layout

The experimental fields were settled at the research farm of Department of Genetics and Plant Breeding of CCS Haryana Agricultural University, Hisar (29.1504° N, 75.7057° E) during the rabi season of 2015-16. A total of 64 faba bean genotypes were used in the present investigation the faba bean genotypes were procured from the different sources as presented in Table 1. All 64 genotypes were sown in a randomized complete block design (RBD) with three replications in a plot of 3 m length, with a row to row spacing of 30 cm and plant to plant distance of 10 cm. All other plant production practices were followed as defined elsewhere^14^.

**Table 1.** A list of sixty-four faba bean genotypes and their source.

| Genotype name | Code | Source |
| --- | --- | --- |
| HB 3 | G1 | CCSHAU |
| HB 18 | G2 | CCSHAU |
| HB 19 | G3 | CCSHAU |
| HB 24 | G4 | CCSHAU |
| HB 28 | G5 | CCSHAU |
| HB 31 | G6 | CCSHAU |
| HB 33 | G7 | CCSHAU |
| HB 38 | G8 | CCSHAU |
| HB 39 | G9 | CCSHAU |
| HB 48 | G10 | CCSHAU |
| HB 62 | G11 | CCSHAU |
| HB 50 | G12 | CCSHAU |
| HB 174 | G13 | CCSHAU |
| HB 176 | G14 | CCSHAU |
| HB 604 | G15 | CCSHAU |
| HB 613 | G16 | CCSHAU |
| AFB 13-1 | G17 | IGKV |
| AFB 13-2 | G18 | IGKV |
| DFB 13-1 | G19 | NBPGR |
| DFB 13-2 | G20 | NBPGR |
| DFB 10-1 | G21 | NBPGR |
| DFB 10-2 | G22 | NBPGR |
| DFB 10-3 | G23 | NBPGR |
| DFB-1 | G24 | NBPGR |
| NDF-4 | G25 | NDUAT |
| NDF-11 | G26 | NDUAT |
| NDF 12-1 | G27 | NDUAT |
| NDF 13-2 | G28 | NDUAT |
| NDF-9 | G29 | NDUAT |
| NDFB-12 | G30 | NDUAT |
| NDFB-13 | G31 | NDUAT |
| NDFB-14 | G32 | NDUAT |
| NDF 1 | G33 | NDUAT |
| NDF 8 | G34 | NDUAT |
| RFB-8 | G35 | Ranchi, India |
| RFB-9 | G36 | Ranchi, India |
| RFB-11 | G37 | Ranchi, India |
| RFB-12 | G38 | Ranchi, India |
| RMDFB-1 | G39 | Ranchi, India |
| RMDFB-2 | G40 | Ranchi, India |
| MLMK-7 | G41 | CCSHAU |
| Pusa sumit | G42 | NBPGR |
| ET 3134 | G43 | Bulgaria |
| ET 2112 | G44 | Bulgaria |
| ET 4101F | G45 | Bulgaria |
| ET 5108 | G46 | Bulgaria |
| EC 628936 | G47 | Bulgaria |
| EC 329612 | G48 | Bulgaria |
| EC 628937 | G49 | Bulgaria |
| EC 591776 | G50 | Bulgaria |
| EC 591755 | G51 | Bulgaria |
| EC 628955 | G52 | Bulgaria |
| EC 243624 | G53 | Bulgaria |
| EC 628939 | G54 | Bulgaria |
| EC 354951 | G55 | Bulgaria |
| EC 598938 | G56 | Bulgaria |
| EC 243770 | G57 | Bulgaria |
| EC 25085 | G58 | Bulgaria |
| EC 10845 | G59 | Bulgaria |
| EC 628962 | G60 | Bulgaria |
| EC 628942 | G61 | Bulgaria |
| EC 628930 | G62 | Bulgaria |
| EC 10719 | G63 | Bulgaria |
| VIKRANT | G64 | CCSHAU |

\*Where CCS Haryana Agricultural University (CCSHAU), Hisar, India; Indira Gandhi Krishi Vishwavidyalaya (IGKV), Raipur, Chhattisgarh, India; National Bureau of Plant Genetic Resources (NBPGR), New Delhi, India; Narendra Deva University of Agriculture and Technology (NDUAT), Faizabad, India.

### Observations Recorded

The following traits were measured as mentioned below.

#### Days to 50% flowering (DF)

The number of days was estimated starting from the date of sowing to the date when 50 percent of the plants in each genotype in each replication came to bloom.

#### Days to maturity (DM)

The days to maturity were counted from the date of sowing to the date of maturity of the plant, i.e. when the 90-95% pods turn black in colour.

#### Plant height (PH)

The plant height in cm at the time of harvest was recorded at the time of maturity.

#### Number of branches per plant (NPB)

The branches were counted on the plant basis at the time harvesting.

#### Number of cluster per plant (CP)

The cluster number was recorded at the time of harvesting on per plant basis.

#### Number of pods per plant (PP)

Pods number was recorded at the time of harvesting.

#### Pod length (PL)

Randomly ten pods from five plants were measured for average pod length in centimetres.

#### Number of seeds per pod (SP)

A random sample of ten pods from five plants threshed and counted to work out the average number of seeds per pod.

#### 100-seed weight (SWT)

Being a large-seeded crop faba bean were determined the one hundred seeds weight (g) using an electrical balance.

#### Seed yield /plant (GY)

All selected individual plants at maturity were threshed separately, and seed weight (g) of the threshed seed from each plant was recorded on electrical balance.

### Data Analysis

All of the statistical analysis is performed with the help of statgraphics Centurion XVIII software package (Warrenton, VA, USA). Further, the relationship between the sixty-four genotypes was determined with the Unweighted Pair Group Method with Arithmetic Mean (UPGMA) method of hierarchical clustering was applied for the grouping of the sixty-four faba bean genotypes used in this study using the ten descriptors using the R platform (R Core Team 2015). The Pearsons linear coefficient correlations were estimated in R using corrplot package in R environment^15^. We also performed the path coefficient analysis maintaining the seed yield as the dependent variable to see the effect of other traits on the seed yield (GY) with the help of R package Lavaan^16^.

## Results and Discussion

The mean sum of squares are presented in Table 2. The mean sum of squares due to genotypes were highly significant for all the characters, thereby indicating the presence of sufficient genetic variability among the genotypes for all the characters studied (*p* <0.05) (Table 1). Whereas, the UPGMA method based clustering produced four main clusters and several subclusters. The genotypes G13, G14 and G16, existed together as the least number of individuals in a subcluster (Figure 1).

**Table 2.** Analysis of variance for seed yield and its component characters in faba bean.

| Characters | Mean sum of squares |  |  |
| --- | --- | --- | --- |
|  | Replication | Genotype | Error |
|  | 2 | 164 | 128 |
| Days to 50% flowering | 118.28 | 422.70** | 26.28 |
| Days to maturity | 491.85 | 390.32** | 38.06 |
| Plant height (cm) | 171.51 | 539.39** | 169.19 |
| Noumber of branches/ plant | 0.71 | 5.39* | 0.43 |
| Number of cluster/plant | 0.50 | 56.93* | 0.96 |
| Number of pods/plant | 6.82 | 497.56** | 40.65 |
| Pod length (cm) | 0.21 | 3.36* | 0.41 |
| Number of seeds/pod | 0.68 | 0.45* | 0.05 |
| 100 seeds weight (gm) | 0.50 | 1,525.2** | 10.93 |
| Seed yield/plant (gm) | 156.54 | 7,954.6** | 16.68 |

\*\*, \* repesemnt significant at 1% and at 5%, respectively.

**Figure 1.**
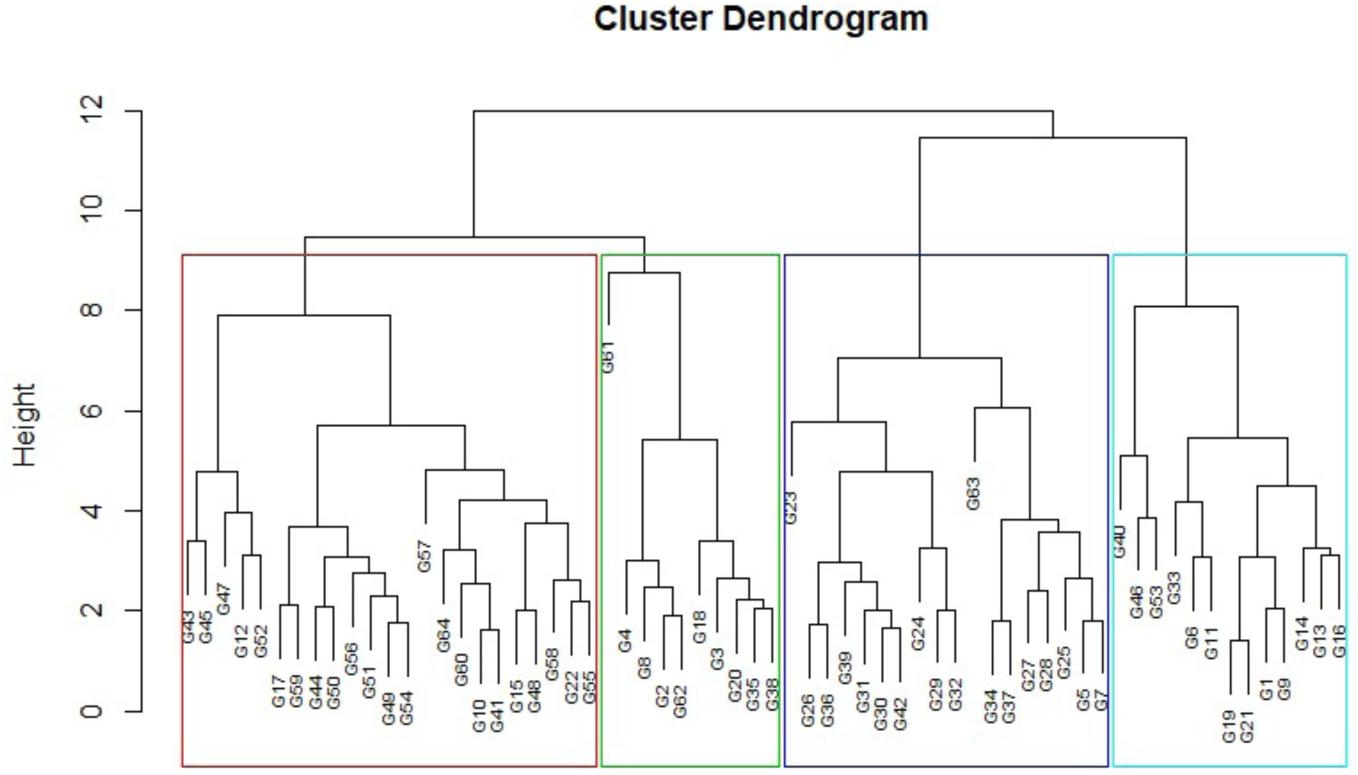
Unweighted Pair Group Method with Arithmetic Mean (UPGMA) clustering of sixty-four faba bean genotypes. The cophenetic correlation coefficient of clustering is 0.7.

### Principal Components Analysis

Plotting the sixty-four genotypes of faba bean via PCA plot revealed a substantial amount of diversity in the faba bean genotypes (Figure 2). Most of the genotypes were cluster together and appear in the centre of the graph. Whereas, the genotypes G24, G40, G52, G61, and G63 appeared as the individuals on the sides of the plots. Pointing out that these genotypes were substantially distinct from the rest of the genotypes (Figure 2).

**Figure 2.**
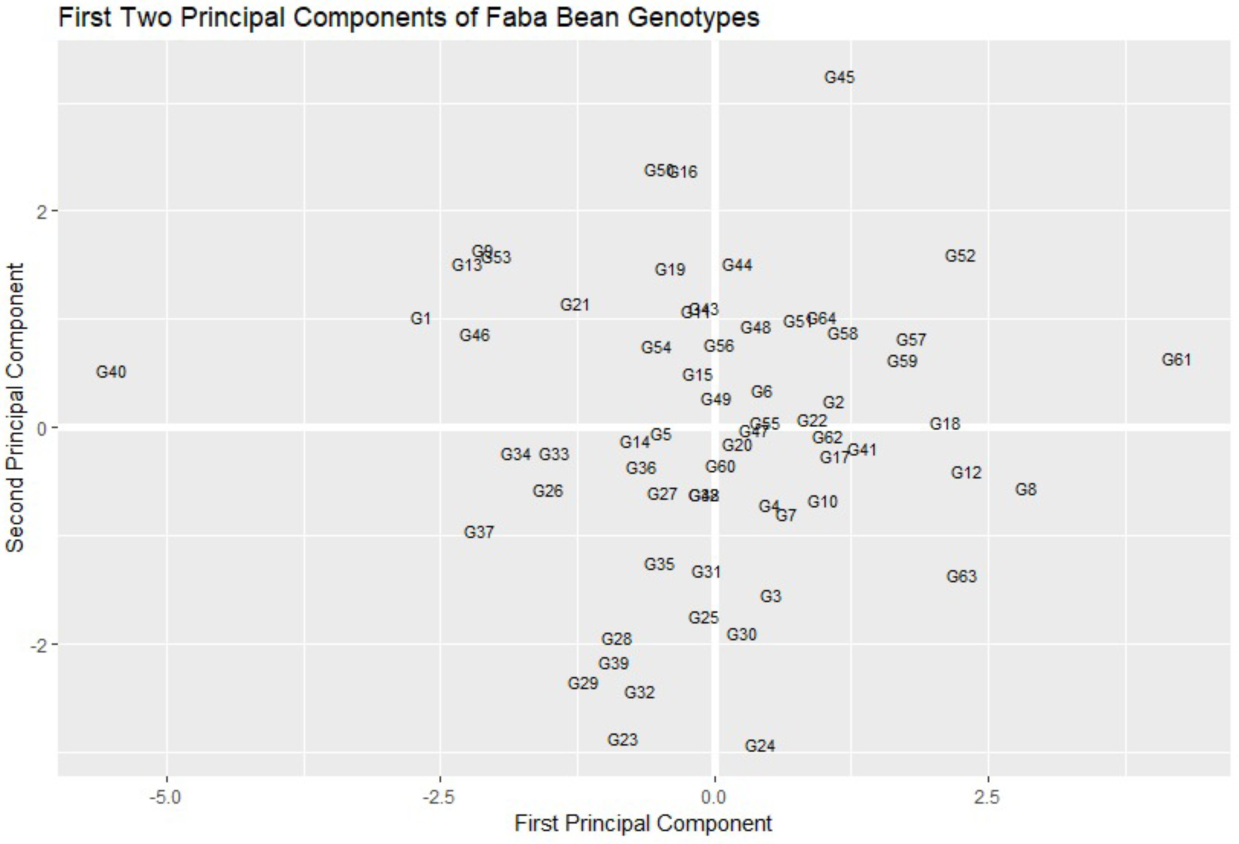
The first (X-axis) and second (Y-axis) principal component projection of sixty-four faba bean genotypes used in the present study.

### Correlation and Path Analysis

The correlation coefficients are represented in Figure 3. The number of cluster per plant (CP) was positively and significantly correlated with the pods per plants (PP) (Figure 3). Whereas, the number of branches per plant (NPB) was positively correlated with the cluster per plant (CP) and also with the number of pods per plant (PP) (Figure 4). Although most of the correlation coefficient values were weak for most of the traits (Figure 3). Therefore, the direct and indirect effect on the seed yield (SY) was determined with the path coefficient analysis is provided in Figure 4. The most substantial positive impact on seed yield was determined by the number of cluster per plant (CP) (Figure 4). In contrast, an adverse effect was determined by the number of pods per plants (Figure 4).

**Figure 3.**
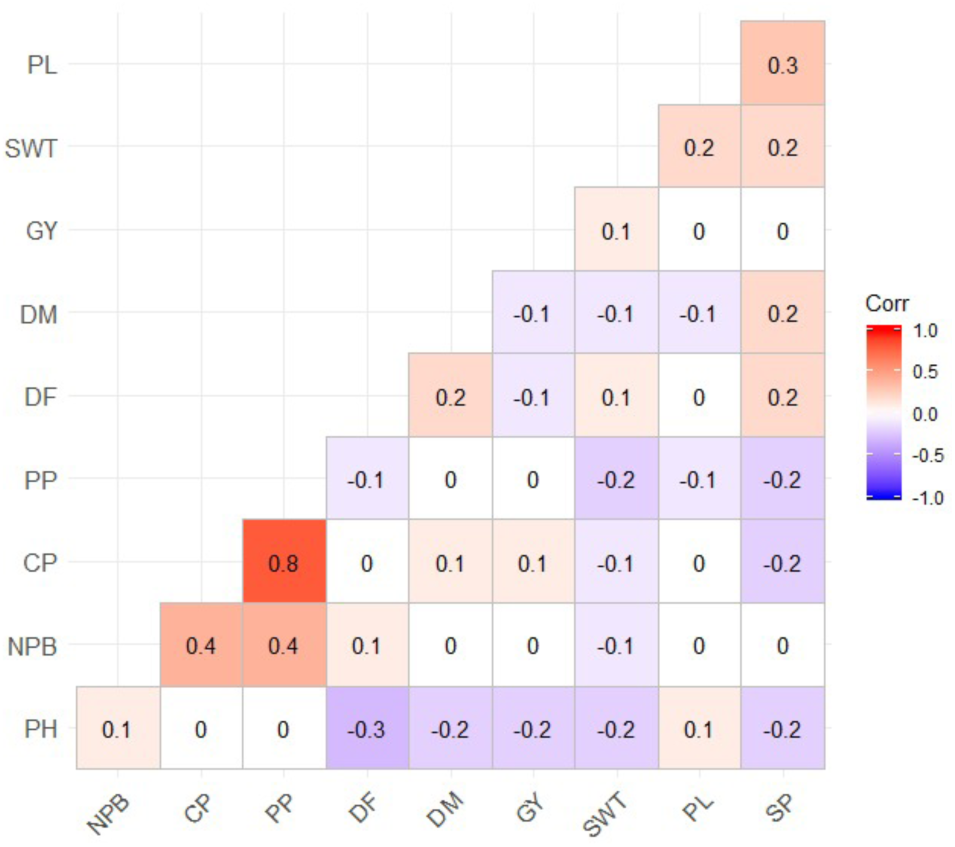
Pearson’s correlation coefficients for the ten traits in the faba bean with significant values *p* < 0.05 highlighted.

**Figure 4.**
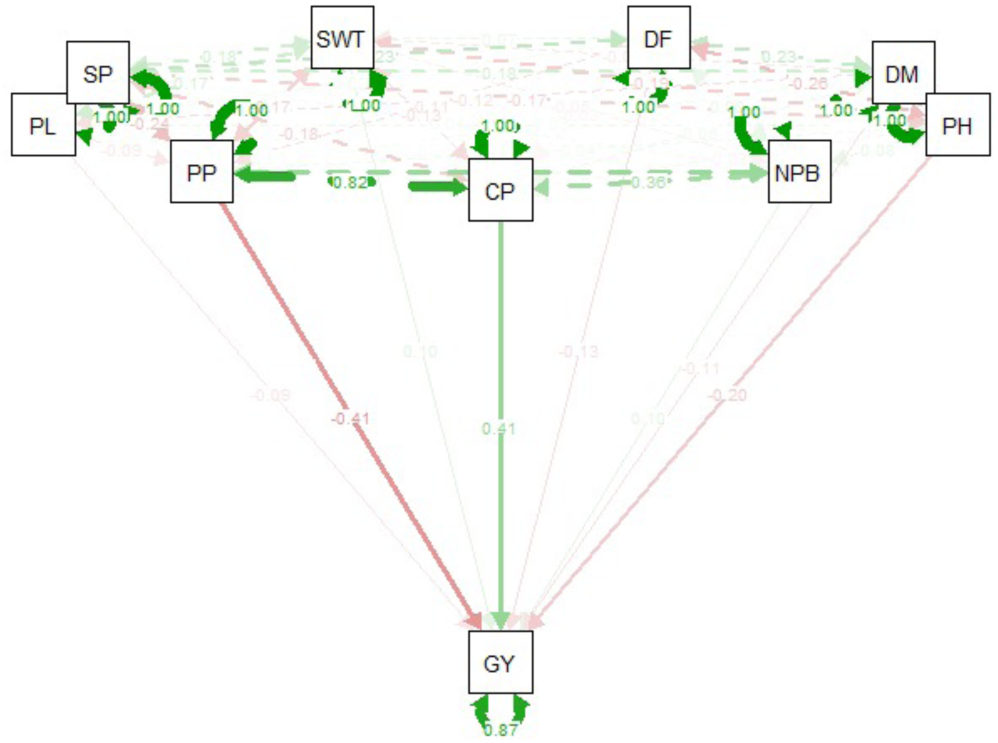
Path coefficient analysis considering the seed yield per plant as the dependent variable on all other remaining traits.

Variability has a unique significance for planning an efficient breeding programme of any crop species. Adequate genetic variability leads to a successful crop improvement programme. Breeder globally collects the germplasm from various agro-climatic regions and evaluate and characterize them for genetic variability. Moreover, both genetic, as well as the environmental factors, contribute to variation among individuals^17,18^. In faba bean Kalia^19^ in 2004 conducted variability and interrelationship studies among twenty-four divergent genotypes. Significant differences were showed with sufficient variability for pod yield and quality characters that could be exploited in breeding. High heritability estimates (97%) along with top genetic advance (126%) for pod yield indicate an additive gene action in its inheritance.

Whereas, Alan and Geren^20^ in 2007 observed the high heritability for seed yield, moderate for seeds per pod and low for pods per plant. These results showed that the environment has a more significant effect on pods per plant and number of stems. Correlation measures the mutual relationship between various plant characters and determines the components on which selection can be based for improvement. The knowledge of correlation that exists between important aspects may facilitate proper interpretation of results and provide a basis for planning more efficient crop improvement programmes^21^. We detected a significant variation for all of the studied traits, and in this direction, Ouji et al.^22^ reported significant variability has present in plant height, petal length and non-significant variability observed in the number of branches in the plant. Therefore information’s regarding the direct and indirect effects of the various components of yield, as obtained from path analysis.

Further, they detected the positive correlation between the number of nodes per plant and number of seeds per plant, date of flowering, number of stem per plant, leaflet length, leaflet width, petal length, petal colour, plant height, pod length, seed length etc. Chaieb et al.^23^ significant positive correlation coefficient was recorded between yield per plant and number of pods per plant (r=0.963) for all studied genotypes. Mustafa^24^ in 2007 revealed that principal component analysis and cluster analysis indicated higher genetic variation within the faba bean in Egypt. Sharifi and Aminpana^25^ in 2014 hierarchical cluster analysis were used for grouping 10 genotypes with 28 traits. Faba bean genotypes were clustered into three main groups. The results of this study showed that days to flowering, days to harvesting, the weight of pod per plant before physiological maturity stage, hundred seed weight and plant height were the traits that closely correlated to dry seed weight per meter square.

## Conclusions

In conclusion, a collection of faba bean (Vicia faba L.) genotypes was evaluated and the results of correlation and path coefficient analysis indicated that the traits like 100-seed weight, number of cluster per plant, number of pods per plant, number of branches per plant and days to flowering should be given due consideration while performing selection for seed yield in segregating generations of faba bean. The genotypes, namely EC-628955, EC-628942, EC-628937, EC-628936, EC-591755, EC-3134, EC-2112, EC-628930, and EC-591776 are selected on the basis of their mean performance for different characters. These genotypes figured importantly to be included in crossing programme to exploit genetic variability among populations further, to affect the selection of superior elite lines for hybridization programme and/or elite populations for composite varieties.

## Significance Improtance

To determine and use the diversity for the faba bean improvement it is essential to study the variability, genetic divergence, correlation and path analysis; therefore sixty-four genotypes of faba bean were used to determine the diversity in the faba bean genotypes.

